# Brain mechanisms underlying the emotion processing bias in treatment-resistant depression

**DOI:** 10.1101/2023.08.26.554837

**Authors:** Xiaoxu Fan, Madaline Mocchi, Bailey Pascuzzi, Jiayang Xiao, Brian A. Metzger, Raissa K. Mathura, Carl Hacker, Joshua A. Adkinson, Eleonora Bartoli, Salma Elhassa, Andrew J. Watrous, Yue Zhang, Wayne Goodman, Nader Pouratian, Kelly R. Bijanki

## Abstract

Depression is associated with a cognitive bias towards negative information and away from positive information. This biased emotion processing may underlie core depression symptoms, including persistent feelings of sadness or low mood and a reduced capacity to experience pleasure. The neural mechanisms responsible for this biased emotion processing remain unknown. Here, we had a unique opportunity to record stereotactic electroencephalography (sEEG) signals in the amygdala and prefrontal cortex (PFC) from 5 treatment-resistant depression (TRD) patients and 12 epilepsy patients (as control) while they participated in an affective bias task in which happy and sad faces were rated. First, compared with the control group, patients with TRD showed increased amygdala responses to sad faces in the early stage (around 300 ms) and decreased amygdala responses to happy faces in the late stage (around 600 ms) following the onset of faces. Further, during the late stage of happy face processing, alpha-band activity in PFC as well as alpha-phase locking between the amygdala and PFC were significantly greater in TRD patients compared to the controls. Second, after deep brain stimulation (DBS) delivered to bilateral subcallosal cingulate (SCC) and ventral capsule/ventral striatum (VC/VS), atypical amygdala and PFC processing of happy faces in TRD patients remitted toward the normative pattern. The increased amygdala activation during the early stage of sad face processing suggests an overactive bottom-up processing system in TRD. Meanwhile, the reduced amygdala response during the late stage of happy face processing could be attributed to inhibition by PFC through alpha-band oscillation, which can be released by DBS in SCC and VC/VS.

## Introduction

Major depressive disorder (MDD) is characterized by excessive low mood and a reduced capacity to experience pleasure. According to Beck’s cognitive model of depression, the biased acquisition and processing of information play a central role in the development and maintenance of depression (Beck 1967; Beck 2008). Numerous studies have shown that individuals with depression tend to have a negative bias across various cognitive domains, including perception (Disner et al., 2011; Roiser et al., 2012), attention (Gotlib et al., 2004), and memory (Hamilton and Gotlib 2008). For example, depressed patients tend to interpret neutral faces as sad and happy faces as less happy (Bourke et al., 2010). Understanding the neural mechanism responsible for the biased processing of emotional stimuli in depression might bring important clinical benefits, including predicting, detecting, and treating depression.

The biased processing of emotional stimuli in depression has been linked to dysfunction in amygdala (Arnone et al., 2012; Young et al., 2017), prefrontal cortex (Siegle et al., 2007; Zhong et al., 2011), dorsal anterior cingulate (dACC) (Greicius et al., 2007; Foland-Ross et al., 2013) and their connections (Ramasubbu et al., 2014; Cheng et al., 2018). Researchers have proposed that cognitive biases in depression are due to maladaptive bottom-up processes which can alter perceptions of the environment and social interactions (Cheng et al., 2016; Clark and Beck 2010). Consistent with this hypothesis, fMRI studies indicate that depressed patients exhibit potentiated amygdala reactivity to sad faces and reduced responsiveness to happy faces, even in the absence of conscious awareness of the picture, suggesting automatic mood-congruent cognitive biases in depression (Suslow et al., 2010; Victor et al., 2010). However, a correlation between this automatic amygdala response and current depression severity was only observed when processing happy faces, but not sad faces (Suslow et al., 2010). One possibility is that there are separate neural mechanisms that increase the salience of negative stimuli and decrease the salience of positive or rewarding stimuli in depression. The reduced responsiveness of the amygdala to happy faces in individuals with depression could be due to the overregulation of the amygdala by the prefrontal cortex (Drevets 2001; Almeida et al., 2010; Cheng et al., 2016). Consistent with this hypothesis, an fMRI study using dynamic causal modeling found increased suppressive influences from orbitomedial prefrontal cortex to amygdala during classification of happy faces in depression (Almeida et al., 2009). But in other fMRI studies, depressed patients showed a wide-spread reduction in the intrinsic connectivity between amygdala and prefrontal cortex (Ramasubbu et al., 2014; Dannlowski et al., 2009). In some studies, researchers even proposed that decreased PFC activity reduces nucleus accumbens and amygdala reactivity in people with depression, which, in turn, contributes to the inability to adaptively alter reward-seeking behavior (Heller et al., 2009; Pizzagalli et al., 2009; Epstein et al., 2006). Due to the predominant use of methods with either high spatial and poor temporal resolution (e.g., fMRI) or high temporal and poor spatial resolution (e.g., EEG/MEG) in previous studies, the neural mechanism responsible for the biased processing of emotional stimuli in depression is far from fully understood. Here we use human intracranial stereo-electroencephalography (sEEG) recordings with high spatial and temporal resolution to investigate the neurobiological mechanisms that contribute to the selective processing towards negative and away from positive stimuli in patients with treatment-resistant depression (TRD).

We recorded sEEG signals simultaneously from amygdala and prefrontal cortex (PFC) in 5 TRD patients and 12 pre-surgical epilepsy patients (as control) while they participated in an affective bias task in which happy and sad faces were rated. Consistent with previous findings, our results revealed increased amygdala responses to sad faces and decreased amygdala responses to happy faces in TRD patients compared to controls. With the help of the excellent spatial and temporal resolution of sEEG, we further observed that increased amygdala responses to sad faces emerged at an early stage (around 300 ms), while decreased amygdala responses to happy faces emerged at a late stage (around 600 ms). Importantly, during this later stage of decreased amygdala responses to happy faces, we found an increased alpha-band activity in PFC, as well as greater alpha-phase locking (connectivity) between the amygdala and PFC in TRD patients compared to the controls. After the delivery of deep brain stimulation (DBS) to two hubs that connect cortical and subcortical network regions relevant to the expression of depressive symptoms, the atypical amygdala and PFC activity, along with their connections while rating happy faces in TRD patients, reverted toward the normative pattern. Thus, our results provide important direct electrophysiological evidence for the neural mechanisms underlying the biased processing of emotional stimuli in depression. The increased amygdala activation during the early stage of rating sad faces suggests an overactive bottom-up processing system. While the reduced amygdala response during the late stage of rating happy faces may be attributed to increased inhibition by PFC through alpha-band oscillation.

## Results

Five TRD patients were implanted with sEEG electrodes for neural recordings, which were used to conduct a thorough search of stimulation parameter space in order to build a comprehensive understanding of the pathophysiology of treatment-resistant depression, as well as the neural responses to stimulation therapy (Sheth et al., 2022). Twelve epilepsy patients were implanted with sEEG electrodes for the purpose of seizure focus mapping. These epilepsy patients are here viewed as a control group, as they represent a range of depressive comorbidities, but they do not have treatment-resistant major depressive disorder. Prior to exploring DBS parameters in TRD patients and performing resection surgery in the control group, we recorded local field potentials (LFPs) from PFC (Total of 180 contacts in TRD and 119 contacts in control) and amygdala (36 contacts in TRD and 52 contacts in control) (**Table 1 and Figure 1B**) while TRD and epilepsy patients participated in the affective bias task in which happy and sad faces are rated (**Figure 1A**).

**Table 1.** Number of contacts in each patient.

|  | Patient | Left |  | Right |  |
| --- | --- | --- | --- | --- | --- |
|  |  | PFC | Amygdala | PFC | Amygdala |
| TRD | Dep1 | 21 | 4 | 26 | 5 |
|  | Dep2 | 19 | 3 | 27 | - |
|  | Dep3 | 16 | 5 | 14 | 4 |
|  | Dep4 | 11 | 4 | 10 | 4 |
|  | Dep5 | 16 | 4 | 20 | 3 |
| Control | EP1 | 9 | 2 | - | - |
|  | EP2 | - | - | 12 | 4 |
|  | EP3 | - | 5 | 8 | - |
|  | EP4 | 8 | 5 | - | - |
|  | EP5 | - | 5 | - | 5 |
|  | EP6 | 9 | 4 | 2 | 4 |
|  | EP7 | 15 | 2 | 9 | - |
|  | EP8 | - | - | 13 | 2 |
|  | EP9 | - | 5 | 11 | - |
|  | EP10 | - | 6 | - | 3 |
|  | EP11 | 5 | - | - | - |
|  | EP12 | 9 | - | 9 | - |

**Figure 1.**
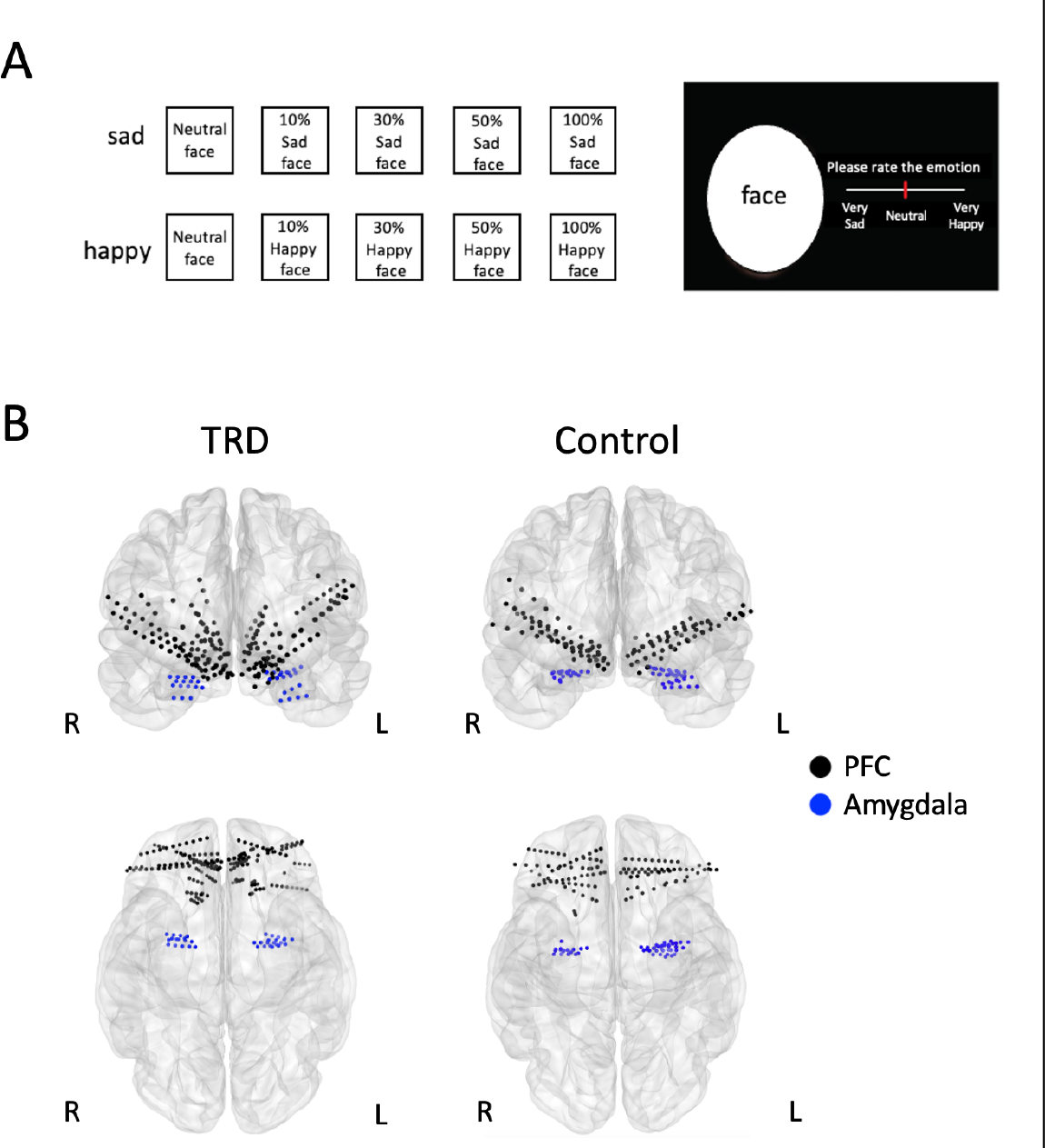
(**A**) Affective bias task. The stimulus set includes morphed faces from maximal emotional intensity to neutral (100% sad, 50% sad, 30% sad, 10% sad, neutral, 10% happy, 30% happy, 50% happy and 100% happy). Participants were asked to rate the emotional intensity of each stimulus presented on the screen by clicking a location on the slider bar. (**B**) Spatial distribution of recording contacts across 5 TRD patients and 12 epilepsy patients in the MNI space. Top: Coronal view. Bottom: Ventral view. Contacts within PFC are black, and electrodes within amygdala are blue. (L, left; R, right). Only contacts in PFC and amygdala are shown.

### Temporal dynamics of amygdala responses in TRD

Given the central role of amygdala in detecting and interpreting emotion information, we first examined intracranial event-related potential (iERP) in amygdala while participants rated sad and happy faces. Left, right and bilateral amygdala iERP response to emotional faces from the two groups are shown in **Figure 2A**. In each group and each condition, there is a large negative potential peaking around 300 ms followed by a small positive potential. We define the duration of the large negative potential as the early stage of amygdala response and the small positive potential as the late stage of amygdala response (**Figure 2B**). In all conditions (sad and happy), the early stage of amygdala response lasts longer in TRD compared to control group. Then, we calculated the peak amplitude of each stage in each group and each condition. We combined results from left and right amygdala because similar results were obtained. Peak amplitude of early stage was significantly larger in TRD compared to control group during sad face processing (**Figure 2C**, TRD: -12.9±1.7, control: -7.5±1.1; t_86_=2.695, p=0.009), but not during rating happy faces (**Figure 2C**, TRD: -11.0±1.3, control: -8.4± 1.1; t_84_=1.567, p=0.121). Moreover, the peak amplitude of late stage was significantly smaller in TRD patients compared to control group in happy face condition, but not in sad face condition (**Figure 2C**, happy faces: TRD: 1.9 ±1.2, control: 7.6±1.0, t_84_=3.610, p=0.001; sad faces: TRD: 7.6±1.0, control: 6.7±1.0, t_86_=0.651, p=0.517). Thus, compared to control group, TRD patients displayed increased amygdala response to sad faces at an early latency, and decreased amygdala response to happy faces at a late latency.

**Figure 2.**
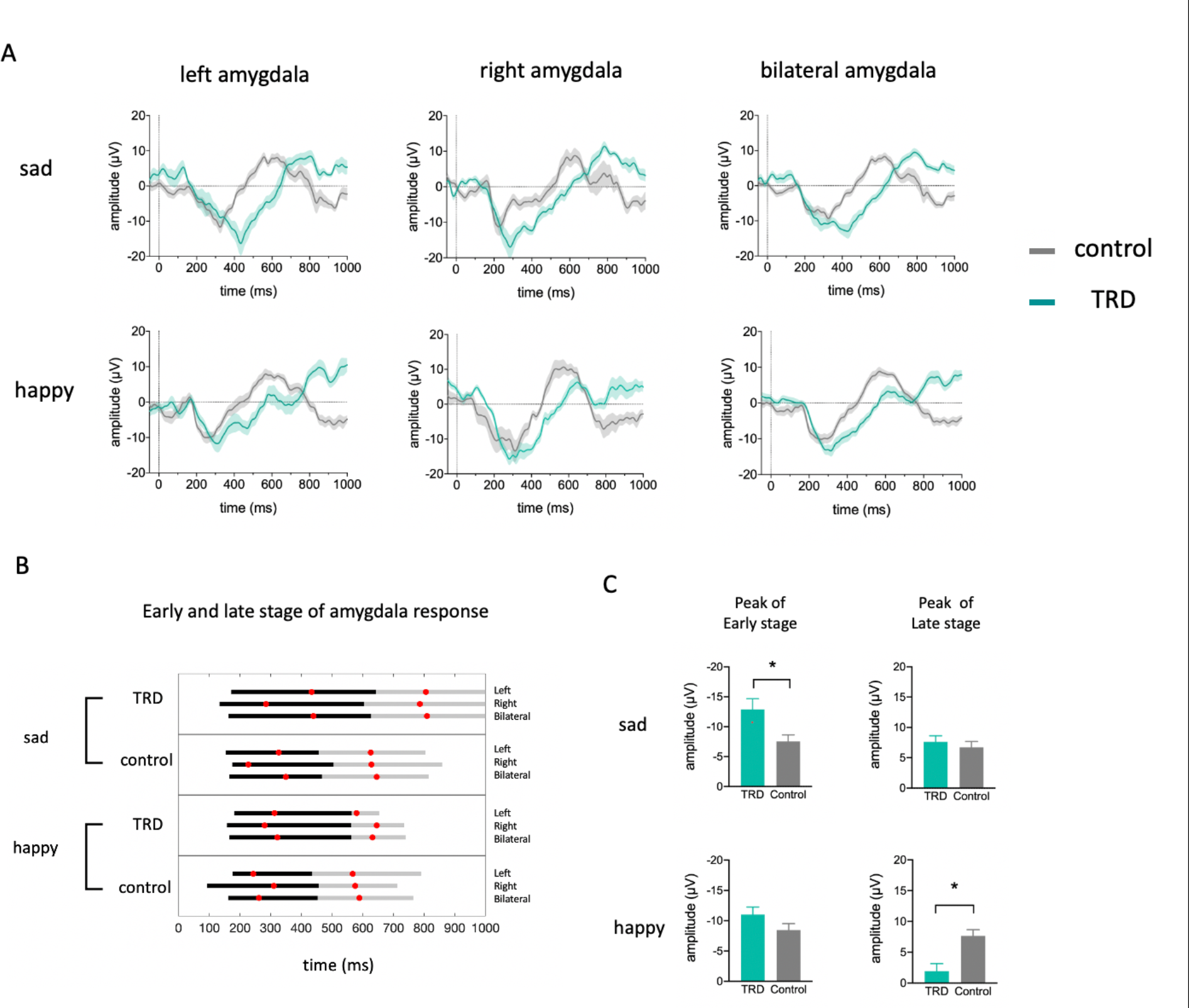
Amygdala iERP response to sad and happy faces. (**A**) Amygdala iERPs from five TRD patients (20 contacts in left amygdala and 16 contacts in right amygdala) and twelve epilepsy patients (34 contacts in left amygdala and 18 contacts in right amygdala) to sad and happy faces were averaged across contacts. Shaded area indicates s.e.m. (**B**) Time windows of early (black horizontal bars) and late stage (gray horizontal bars) amygdala iERP response. Red dot depicts the time point of averaged peak amplitude. (**C**) Peak amplitude differences between TRD and control within early stage and late stage for each condition (sad and happy faces). * p<0.05 (unpaired t tests).

### TRD exhibit greater alpha-power in PFC at the late stage

The altered amygdala response we observed in TRD patients could be due to the regulation from prefrontal cortex. Here, we examined whether alpha power (8-12 Hz) in PFC, which may reflect excessive inhibitory processes (Neuper et al., 2001; Engel et al., 2001; Foxe and Synder 2011; Jensen and Mazaheri 2010; Klimesch et al., 2007), is enhanced or reduced in TRD at early or late stage. A three-way ANOVA revealed significant main effects of emotion category (sad vs happy, F_(1,297)_=17.67, p<0.001), time window (early vs late, F_(1,297)_=29.51, p<0.0001) and patient group (TRD vs control, F_(1,297)_=8.230, p=0.0044). The significant interactions of time window by patient group (F_(1,297)_=16.98, p<0.0001) and emotion category (F_(1,297)_=5.993, p=0.0149) indicated that the amount of information about each of these dimensions differs across time. Subsequently, the Sidak’s multiple comparisons test showed significant differences between TRD and control group at the late stage of happy face (adjusted p<0.0001) and sad face (adjusted p=0.0325) processing (**Figure 3A**). In summary, our results showed that there is no difference in PFC alpha-band power between TRD and control group at the early stage. However, at the late stage, TRD patients displayed significantly greater alpha power in PFC than control group, no matter which emotional faces (sad or happy) were processed. The distribution of alpha power at the late stage in each group was shown in **Figure 3B**.

**Figure 3.**
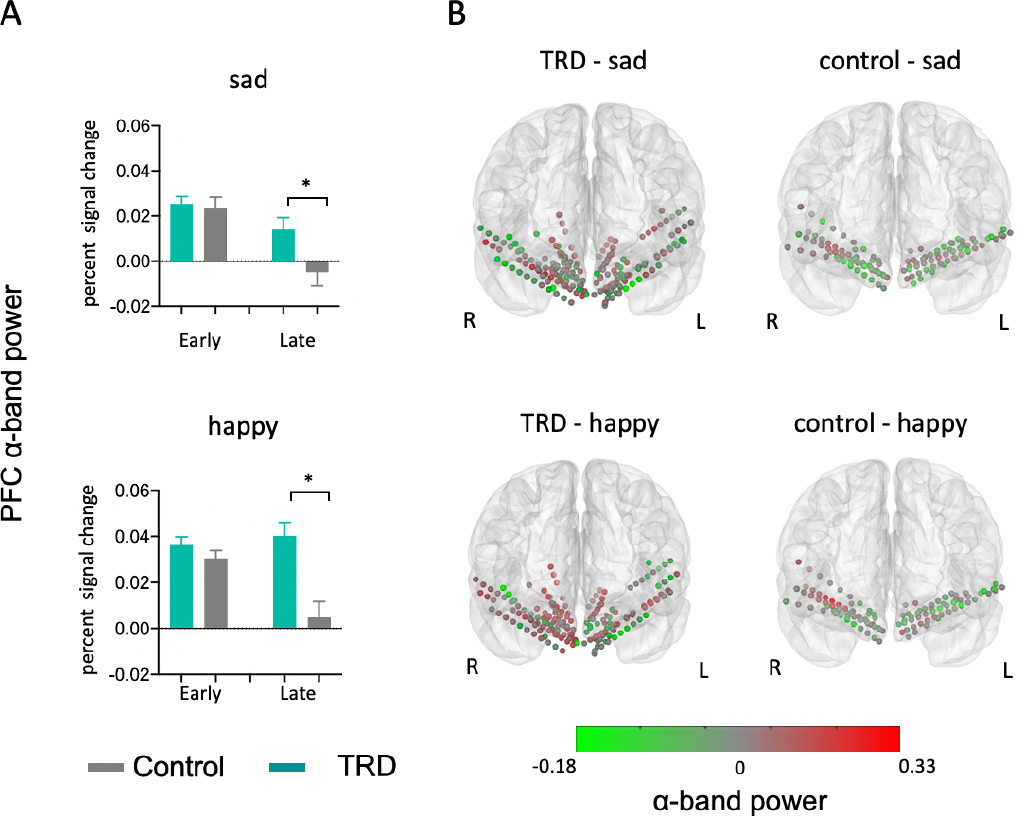
(**A**) Averaged alpha-band power in PFC across 180 contacts in TRD and 119 contacts in control group. * means adjusted p<0.05. (**B**) Coronal view of the alpha-band power (percent signal change) of each contacts within PFC for each group and each condition in the time window of late stage (control-sad: 467-815 ms; control-happy: 453-765 ms; TRD-sad:627-1000 ms; TRD-happy: 563-740 ms). Each circle represents a single contact.

### Increased alpha amygdala-PFC synchrony when TRD patients process happy faces

Given the increased PFC alpha power and decreased amygdala iERP response at the late stage of rating happy faces in TRD, we further hypothesized that the inhibition from PFC to amygdala is increased in TRD at the late stage of processing happy faces. To study this hypothesis, we measured changes in the phase-locking value (PLV) of alpha oscillations between amygdala and PFC during sad and happy face processing, which is a measure of connectivity between two regions. Only a subset of patients (n_TRD_=5, n_control_=8) with at least one contact in both amygdala and PFC were involved in this data analysis. TRD showed greater PLV during happy face trials than control from 608 to 792 ms (**Figure 4A**.Cluster-based permutation test with a cluster threshold P < 0.05). No time cluster expressing a significant group difference was observed in sad face condition (**Figure 4A**). In addition, we calculated the mean PLV within the late stage of happy face processing, and results showed that alpha-band PLV between PFC and amygdala is significantly higher in TRD than control group (**Figure 4B**, unpaired t test, t_11_=3.953 p=0.0023).

**Figure 4.**
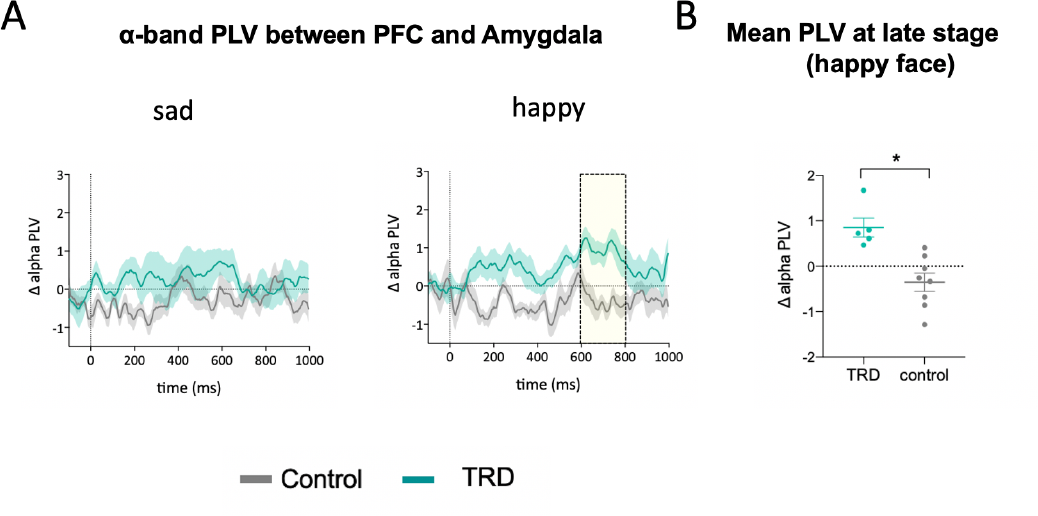
(**A**) Average time courses (mean ± SEM) of alpha PLV changes from baseline between TRD (5 patients) and control group (8 patients). Yellow shade area indicates the time windows in which responses to each stimulus type (sad and happy faces) are significantly different between TRD and control group with a cluster threshold of p < 0.05. (**B**) Mean PLV within the late stage time window of amygdala response to happy faces. Each dot means a single participant. * p<0.05.

In summary, our results revealed separate neural mechanisms for the biased processing of emotional stimuli in TRD (**Figure 5**). When individuals with TRD process sad faces, they show amygdala reactivity that is more intense at the early stage and longer lasting than in healthy controls. While processing happy faces, TRD patients display decreased amygdala response, increased alpha power in PFC and enhanced alpha synchrony between these two regions at the late stage, compared to control group.

**Figure 5.**
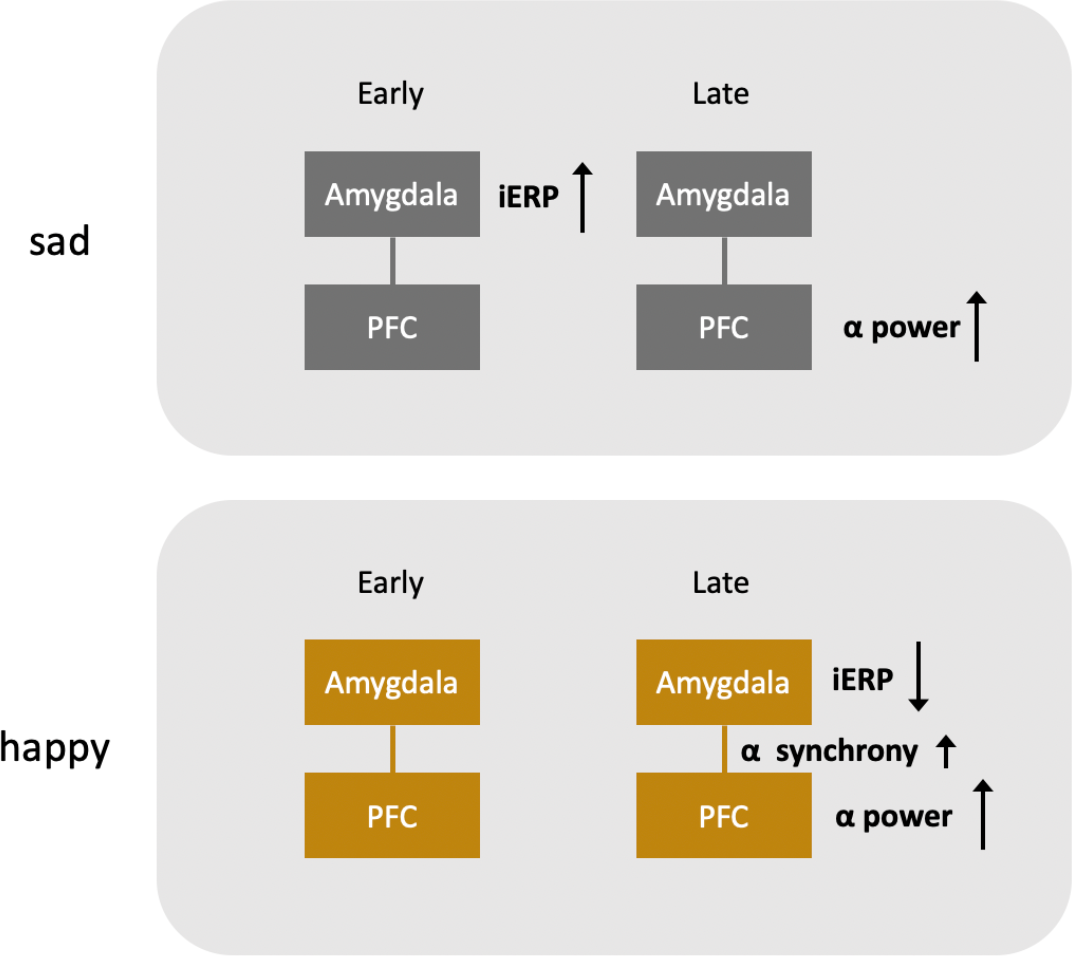
Summary of the results in TRD patients. Arrows indicate higher or lower signal in TRD than control group.

### DBS altered the neural responses to emotional stimuli in TRD

All five TRD patients were asked to perform the affective bias task again after receiving DBS in the subcallosal cingulate (SCC) and the ventral capsule/ventral striatum (VC/VS) (**Figure 6A**). We recorded and analyzed LFPs from sEEG contacts in PFC and amygdala (**Figure 1A**) as before. First, amygdala iERP responses to sad and happy faces were significantly increased after DBS in a late time window (**Figure 6B** left column, sad: 563-713 ms; happy: 598-796 ms, cluster-based permutation test with cluster threshold p < 0.05). The shape of the early-stage amygdala iERP response was not changed by DBS. Second, the alpha-band power in PFC was significantly reduced after DBS in both early and late stage when TRD patients process happy faces (**Figure 6C**, TRD pre-DBS happy early vs TRD post-DBS happy early: adjusted p< 0.001; TRD pre-DBS happy late vs TRD post-DBS happy late: adjusted p< 0.001). Interestingly, TRD patients did not display changed alpha-band power in PFC at any stage in sad face condition (**Figure 6C**, TRD pre-DBS sad early vs TRD post-DBS sad early: adjusted p=0.993; TRD pre-DBS sad late vs TRD post-DBS sad late: adjusted p=0.9474). Third, in the happy face condition, the PLV is reduced to an intermediate pattern between TRD pre-DBS and control group. Overall, DBS in SCC and VC/VS reversed the atypical activity in amygdala and PFC as well as their connectivity toward the normative pattern, so that TRD patients processed happy faces more similarly to controls.

**Figure 6.**
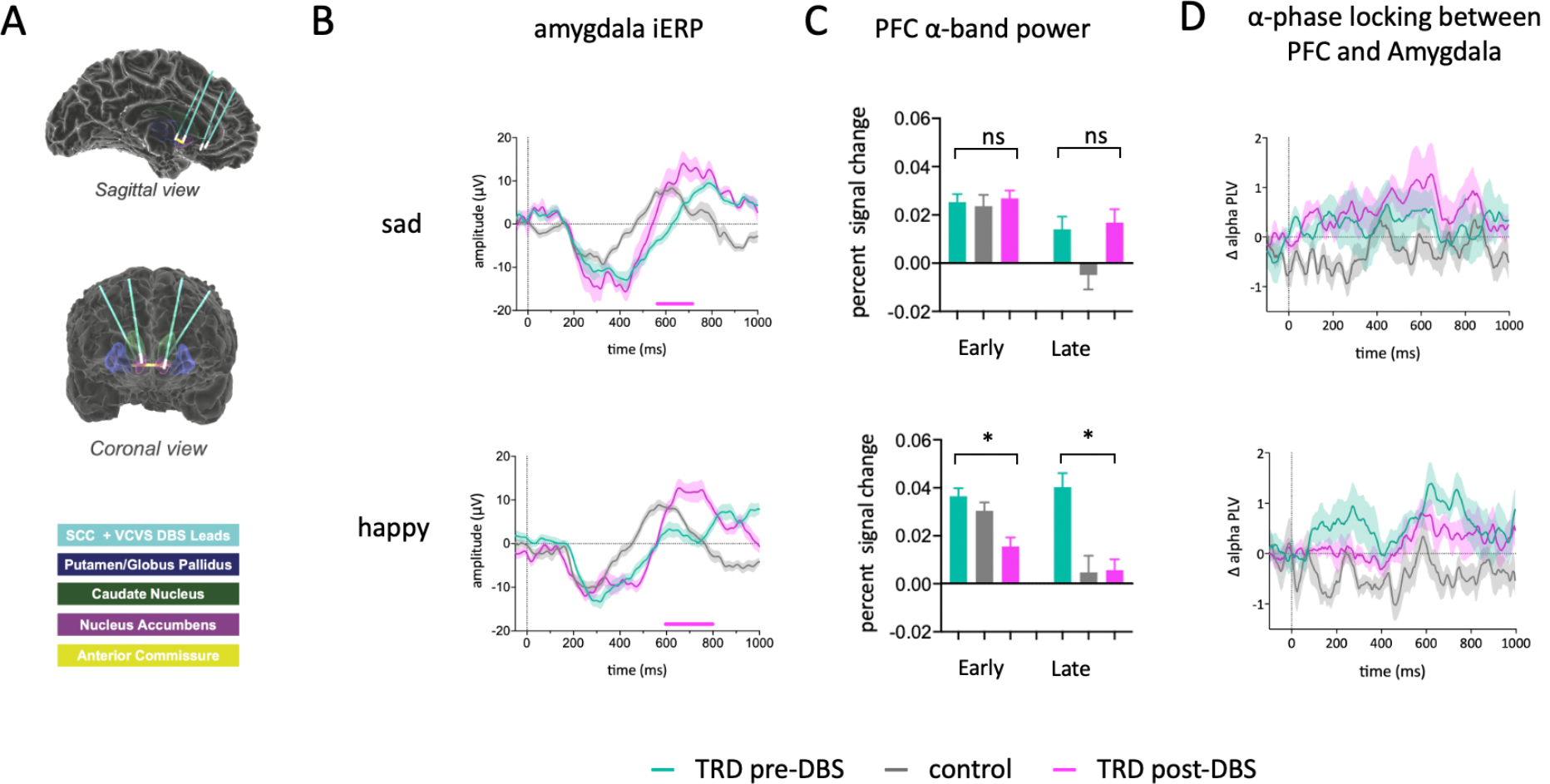
(**A**) Schematic of DBS leads placement in 5 TRD patients. (**B-D**) Data from control group were shown again here as a reference (gray), but not included in statistical analysis. (**B**) Amygdala iERP responses to sad and happy faces were averaged across contacts for each group (TRD pre-DBS, control and TRD post-DBS). Pink horizontal bar below means significant difference between TRD pre-DBS and TRD post-DBS. (**C**) Averaged alpha-band power in PFC across contacts. A three-way ANOVA revealed significant main effects of DBS (pre vs post, F_(1,179)_=20.14, p<0.0001), significant interactions of DBS effect by emotion category (F_(1,179)_=26.69, p<0.0001) and stage (F_(1,179)_=9.209, p<0.0028). The Sidak’s multiple comparisons test showed significant differences between TRD-pre DBS and TRD-post DBS at the early (adjusted p<0.0001) and late stage (adjusted p<0.0001) of happy face processing. * means adjusted p<0.05. (**D**) Averaged time courses (mean ± SEM) of alpha PLV changes from baseline across participants (TRD pre-DBS, control and TRD post-DBS).

## Method

### Participants

Data were obtained from twelve epilepsy patients (7 Female, 5 Male, Age 19-57) and five treatment-resistant depression (TRD) patients (3 Female, 2 Male, Age 32-56). Twelve epilepsy patients were undergoing stereoelectroencephalography (sEEG) monitoring at Baylor College of Medicine for seizure localization before surgical resection. All epilepsy participants completed questionnaires on depressive symptoms (Beck Depression Inventory, BDI, Beck et al., 1987). The average BDI score is 21.17±11.7 (mean±SD). Participants with TRD involved in this study (N =5) were enrolled in an early feasibility trial (ClinicalTrials.gov Identifier: NCT03437928) of individualized deep brain stimulation guided by intracranial recordings (Sheth et al., 2022). Each TRD patients was implanted with ten temporary sEEG electrodes for neural recordings and four permanent deep brain stimulation leads for stimulation delivery (sEEG electrodes are shown in **Figure 1B**, DBS leads are shown in **Figure 6A**). TRD study inclusion criteria include failure of conventional therapies (pharmacological, cognitive/behavioral, electroconvulsive), severity of symptoms, and the ability to provide informed consent. Participants with history of psychosis, personality disorder, recent suicide attempt, or neurodegenerative disorders were excluded from recruitment. This clinical trial is funded by the NIH Brain Research Through Advancing Innovative Neurotechnologies (BRAIN) Initiative (UH3 NS103549) and approved by the FDA (IDE number G180300). All the experiments in this paper involved epilepsy patients and TRD patients were approved by the Institutional Review Board at Baylor College of Medicine (IRB: H-43036, IRB: H-18112). Written informed consent was obtained from each participant.

### Electrode Implantation and localization

DBS leads (Cartesia, Boston Scientific) and sEEG electrodes (Depthalon, PMT Corporation) were implanted using a Robotic Surgical Assistant (ROSA, Zimmer Biomet). The locations of sEEG electrodes in epilepsy patients were entirely based on medical considerations (detection of the seizure foci). All epilepsy patients involved in our study have at least one electrode in PFC or amygdala. Each TRD patient had 10 sEEG electrodes (5 per hemisphere in the prefrontal and mesial temporal regions) and 4 DBS leads (VC/VS and SCC bilaterally). The surgery procedure and DBS leads information can be found in a published study (Sheth et al., 2022). Following surgical implantation, electrodes were localized by co-registration of pre-surgery anatomical T1 MRI scans and post-implantation CT scans using FreeSurfer (https://surfer.nmr.mgh.harvard.edu). Electrode positions were manually marked using BioImage Suite (Papademetris et al). iELVis (Groppe et al., 2017) was used to overlay electrode location into the MRI. Electrodes were then assigned to PFC and amygdala by independent expert visual inspection. We projected electrode positions onto MNI space and displayed on the cortical surface of FreeSurfer ‘fsaverage’ brain for visualization in **Figure 1B** and **Figure 6A**.

### Affective Bias Task

Participants in our study were asked to rate emotional human face photographs, which were displayed on a Viewsonic VP150 monitor with a resolution of 1920 x 1080, positioned at a distance of 57 cm from the participants. Happy, sad and neutral face exemplars (6 identities each; 3 male, 3 female) adapted from the NimStim Face Stimulus Set (Tottenham et al., 2009) were morphed using a Delaunay tessellation matrix to generate subtle facial expressions ranging in emotional intensity from neutral to maximally expressive in steps of 10%, 30%, 50%, and 100% for happy and sad faces, respectively (**Figure 1A**). The experiment was programmed in MATLAB, using Psychtoolbox (Brainard 1997).

In each trial, a white fixation cross was displayed on a black background for 1000 ms (jittered +/-100 ms) and then a face and a rating prompt appeared simultaneously on the screen, positioned on the left and right sides, respectively (as illustrated in **Figure 1A**). Participants were asked to indicate their rating by clicking a specific location on the slider bar using a computer mouse. The ratings were recorded using a continuous scale that ranged from 0 (‘Very Sad’) to 0.5 (‘Neutral’) to 1 (‘Very Happy’). Stimuli were presented in a blocked design in which all happy faces (plus neutral) appeared in one block while all sad faces (plus neutral) appeared in a separate block. There were 30 trials in each block (6 trials for each intensity level). Participants completed three alternating happy and sad blocks. The order of happy and sad blocks was counterbalanced across participants. TRD patients undergo sEEG research for ten days in the EMU; they performed the affective bias task twice after surgical implantation, at day 1 or day 2 (before DBS parameter exploration) and day 8 or day 9 (after DBS parameter exploration). Epilepsy patients completed the task during their in-patient stay (at least 2 hours after any seizure activity was detected).

### Data acquisition and preprocessing

Intracranial EEG data were recorded during affective bias task using Cerebus data acquisition system (Blackrock Neurotech). All neural signals were recorded at a 2000-Hz sampling rate (online band-pass filter 0.3-500 Hz) with a 256-channel amplifier and referenced to a contact in the white matter. Some runs of patient DBSTRD010 were collected at 30-kHz for reasons pertaining to other studies and were downsampled to 2000-Hz. A photodiode was laced in the lower right-hand corner of the screen to mark the trial onsets and its analog voltage response was recorded by the data acquisition system to ensure precise synchronization. Electrode contacts and epochs contaminated with excessive artifacts and epileptiform activity were removed from data analysis by visual inspection. After that, the continuous iEEG data in each electrode contact was notch-filtered at 60 Hz, re-referenced to the common average reference, and segmented from −500 to 1000 ms relative to stimulus onset. Neutral face epochs in each block were removed from data analysis.

### Quantification and statistical analysis

#### Event-related potentials

The segmented data were band-pass filtered from 1 to 30 Hz using a finite impulse response filter (FIR, MNE-Python, Gramfort et al. 2013). iERPs were calculated for each condition (happy and sad), each group (TRD, control) and each contact by averaging the filtered epochs and normalizing them to the mean signal of baseline period (-500 to 0 ms relative to stimulus onset). The iERP time courses were averaged across contacts within left amygdala, right amygdala and bilateral amygdala for each condition (**Figure 2A**). The data from two contacts in patient Dep3 in happy blocks was excluded from time course and amplitude analysis as an extreme outlier (> 8 SDs away from mean). Based on the averaged amygdala iERP waveform across contacts, we define the duration of the large negative potential as the early stage of amygdala response and the small positive potential as the late stage of amygdala response. Specifically, the duration of the early and late stage of amygdala response are defined as same polarity time points surrounding the peak of negative potential and positive potential, respectively (**Figure 2B**). Peak amplitude of each stage was calculated as the averaged amplitude within 150 ms time window of peak (**Figure 2C**). Group differences in peak amplitude were tested with an unpaired t-test.

#### Power analysis

For each recording contact, alpha-band power was estimated for early stage and late stage. We band-pass filtered the data (8-12 Hz, IIR filter), applied the Hilbert transform to extract analytic amplitude envelope, squared the results and normalized them by calculating percent change from pre-stimulus baseline (-500 to 0 ms). Results shown in **Figure 3A** were averaged across all the contacts for each group and each condition. We performed three-way ANOVA (emotion category×time window×patient group) on the alpha power data. The Sidak’s multiple comparisons test was used to estimate the differences between TRD and control group and the effects were reported using adjusted *p*<0.05.

#### Inter-regional phase synchrony

Phase-locking value (Lachaux et al., 1999) was calculated for each contact pair between PFC and amygdala. Only a subset of patients (n_TRD_=5, n_control_=8) with at least one contact in both amygdala and PFC were involved in data analysis. The preprocessed data were alpha-band filtered (8–12 Hz) using a FIR filter (order = 4 cycles of the desired signal). Then Hilbert transform was applied and PLV was calculated for each contact pair and normalized to baseline (−500 to 0 ms). Results shown in **Figure 4A** were averaged across participants. For each condition (sad and happy), we statistically compared the PLVs between TRD and control group using a cluster-based permutation test (N = 10,000, cluster P < 0.05, Bonferroni corrected). We also compared the the mean PLV of happy face trials during the late stage (TRD: from 563 to 740 ms; control: from 453 to 765 ms) between TRD and control group (unpaired t-test).

### DBS stimulation

The DBS leads were positioned to target region (SCC and VC/VS) using previously described methodology based on patient-specific diffusion-weighted imaging data (Tsolaki et al., 2017). DBS parameter exploration was initiated 3 days after surgery in TRD patients as described in clinical protocol (FDA IDE number G180300). We delivered stimulation to the target regions using different parameter sets (Cerestim, Blackrock Microsystems) from day 3 to day 9. The stimulation was off when TRD patients were performing affective bias task at day 8 or day 9.

## Discussion

Our results provide important direct electrophysiological evidence for the mechanisms underlying the biased processing of emotional information in depression. During the early stage of rating sad faces, TRD patients displayed an increase in amygdala activation, indicative of an overactive bottom-up processing system to negative emotions. During the late stage of rating happy faces, we found a reduced amygdala response paired with increased PFC-amygdala connectivity, suggestive of an inhibition by PFC through alpha-band oscillation. Furthermore, after the stimulation of SCC and VC/VS, the neural processing of happy faces in TRD patients remitted toward the normative pattern, implying that DBS releases the inhibition from PFC to amygdala during positive information processing. Altogether, our results suggest that separate neural mechanisms are responsible for the biased negative and positive affective information processing in TRD.

The amygdala plays a critical role in emotion processing and response. It interacts with a variety of cortical and subcortical areas, which together evaluate the salience of sensory stimuli and modify the response of the amygdala (Sander et al., 2003). Numerous fMRI studies have shown that individuals with depression exhibit increased amygdala response to sad faces and decreased amygdala response to happy faces compared to healthy controls (Arnone et al., 2012; Victor et al., 2010; Surguladze et al., 2005). Our iERP results clearly showed that TRD patients displayed heightened and prolonged amygdala activity in response to sad faces, compared to control group. More interestingly, the iERP traces from TRD and control patients are almost overlapping before 200 ms, suggesting no difference in the initial response to sad faces in amygdala. This finding aligns with the theory that depressed and non-depressed individuals do not primarily differ in their initial response to negative events but in their ability to recover from the ensuing negative affect (Teasdale and Dent 1987). Additionally, we also observed prolonged activity in the amygdala during the early stage of happy face processing, although without a significant change in amplitude compared to control. Overall, the longer-lasting amygdala response observed in our study could be related to the extended elaborative processing of emotional information in depression (Siegle et al., 2002).

The increased amygdala iERP we observed at the early stage during sad face processing in TRD supports the hypothesis that increased amygdala activity creates a bottom-up signal that biases negative information processing in higher cortical areas and can maladaptively alter perceptions of the environment (Victor et al., 2010). The altered perception of negative information has been associated with decreased cognitive control from DLPFC (Fales et al., 2008; Drevets et al., 2001; Ochsner and Gross 2005). Another possibility could be that the reduced connectivity between the thalamus and dACC leads to a higher flow of information through the subgenual cingulate cortex. This, in turn, heightens the emotional impact of incoming stimuli for individuals with depression (Greicius et al., 2007; Disner et al., 2011). As individuals with depression often show an attentional bias for sad stimuli (Gotlib et al.,2004), it is still unclear how the increased amygdala response to negative stimuli is related to the inefficient attentional disengagement from negative stimuli.

In the current study, TRD patients display decreased amygdala response, increased alpha power in PFC and enhanced alpha synchrony between these two regions at the late stage of happy face processing, compared to control group. Thus, the reduced amygdala response to happy face in TRD could be due to the inhibition from PFC. Consistent with our findings, a fMRI study examined the the functional connectivity between amygdala and orbitomedial prefrontal cortex using dynamic causal modeling, identified increased negative left-sided top-down orbitomedial prefrontal cortex–amygdala effective connectivity in response to happy faces (Almeida et al., 2009). The connectivity results suggest increased inhibition of the left amygdala by left orbitomedial prefrontal cortex in response to positive emotional stimuli. In addition, increased alpha power in the PFC of individuals with depression has been reported in previous EEG studies (Jaworska et al., 2012; Fingelkurts et al., 2007; Shim et al., 2018). This heightened alpha activity may reflect excessive inhibitory processes, which could contribute to the cognitive and emotional symptoms associated with depression. Consistent with this idea, a clinical improvement after antidepressant treatments was also found to be associated with a decrease in PFC alpha activity (Ulrich et al., 1984; Olbrich and Arns 2013).

With the help of high temporal and spatial resolution of iEEG, our results support that separate neural mechanisms are responsible for the biased negative and positive affective information processing in depression. The observed effect of DBS also provides evidence for this hypothesis. Specifically, after DBS in SCC and VC/VS, both the alpha power in PFC and alpha PLV between PFC and amygdala during happy face processing were reduced. However, DBS has little effect on PFC alpha power and alpha PLV between PFC and amygdala during sad face processing. Consistent with our results, decreased positive emotion rather than exaggerated responses to negative stimuli is thought to be a distinctive feature of depression. For example, amygdala activity during happy face processing or positive recall was negatively correlated with current depression severity (Young et al., 2016; Suslow et al., 2010). But no significant correlation between amygdala activity during sad face processing and current depression severity was observed in those studies. Also, it is worth noting that negative processing bias also exist in high-risk populations of depression and individuals suffering from anxiety (Young et al., 2016, Teachman et al. 2012). Similarly, some studies proposed that amygdala hyperactivity during negative autobiographical recall is a trait-like marker of depression, while amygdala hypoactivity during positive autobiographical recall is a state marker that emerges during active disease and returns to normal with symptom remission (Young et al., 2016). Another two studies by our group also showed that the correlation between the affective bias score and depression severity was primarily driven by positive trials in individuals with treatment-resistant depression (unpublished results). However, in people with varying levels of depression, but not TRD, this correlation was predominantly driven by negative trials (unpublished results).

Converging evidence suggests an important role of amygdala in the recovery from MDD. In some studies, individuals with MDD have been trained to upregulate their amygdala hemodynamic response during real-time fMRI neurofeedback training (Young et al., 2018; Young et al.,2017a; Young et al., 2017b). After training, the amygdala hemodynamic response to positive memories or happy faces was increased and depressive symptoms were reduced, suggesting a recovery from depression. In patients with MDD after sertraline treatment, hyperactivation of the amygdala to masked sad faces decreased and hypoactivation of the amygdala to masked happy faces increased (Victor et al., 2010). Consistent with these findings, our results also showed that the processing bias of happy faces reversed toward the normative pattern after DBS. Together with previous findings, the decreased activation of the amygdala to positive stimuli may indicate clinical significance and some antidepressant drugs or cognitive therapies exert their treatment effect by normalizing this emotional processing.

Intracranial recordings in TRD patients before and after DBS parameter exploration allowed this extraordinary opportunity to study the neural mechanism of biased processing of emotional stimuli in depression, but some limitations must be acknowledged. For example, we use epilepsy patients as control group in our study because it is impossible to get sEEG data from healthy humans. Epilepsy patients often display a range of depressive comorbidities, however none of the patients included in this study was diagnosed with treatment-resistant major depressive disorder, thus providing a valid control for the psychiatric condition of interest. Additionally, the spatial coverage of electrodes in TRD patients in the frontal lobe is limited with respect to the epilepsy cohort due to differences in the surgical targets, driven uniquely by clinical purposes. As a consequence, our ability to resolve the contribution of different PFC subdivisions was not possible in the current study, and further evidence is needed to pinpoint the exact source of the PFC inhibitory effects. Nonetheless, our results investigated the functional profile of amygdala, PFC and their connectivity in affective processing and provide direct electrophysiological evidence with high spatial and temporal resolution to understand the critical framework for the biased acquisition and processing of information, which has a primary role in the development and maintenance of depression.

## Acknowledgement

We thank all patients for their participation and all clinical technicians in EMU for providing support during the research recordings. We would like to thank Sheraz Pasha and Victoria Pirtle for their help in data collection in EMU. We would like to thank Christopher Kovach for his assistance in developing the task. This work was supported by fundings from United States National Institutes of Health (R01-MH127006, NIH K01-MH116364 and NIH UH3-NS103549).

## Author contributions

Writing, X.F., E.B., A.J.W., K.R.B., M.M.

Review and Editing, All Authors.

Data analysis, X.F., Y.Z., R.K.M., S.E.

Methodology, J.A.A., B.A.M., K.R.B., N.P.

Conceptualization, X.F., K.R.B., B.A.M., W.K.G.

Funding acquisition, K.R.B., W.K.G., N.P.

Data collection, B.A.M., M.M., B.P., J.X., C.H.

